# From green to red: experimental evidence for pigment-driven snow darkening

**DOI:** 10.64898/2026.08.16.745148

**Authors:** Pablo Almela, Trinity L. Hamilton

## Abstract

Snow algae are major biological drivers of snow darkening in polar and high-alpine environments. However, the direct contribution of algal pigmentation to snow reflectance has remained difficult to quantify because field observations cannot disentangle the effects of pigmentation from variation in biomass, species composition, and snow physical properties. Here, we characterized the optical effects of pigmentation using hyperspectral spectroradiometry to compare green, orange, and red cyst-like cells of a snow-derived *Haematococcus* isolate while controlling for developmental stage and cell abundance. Cysts became more red with increasing astaxanthin concentrations while chlorophyll-a concentrations remained relatively constant. Relative to green cysts, mean reflectance decreased by approximately 30% in orange cysts and 40% in red cysts. Integrated reflectance across the visible spectrum (350–800 nm) was negatively correlated with astaxanthin concentration. These results provide direct experimental evidence that algal pigmentation alone substantially reduces reflectance after controlling for cell abundance and developmental stage, and indicate that differences in snow physical properties may partly obscure this effect under natural field conditions. Our findings identify astaxanthin accumulation as an intrinsic driver of biological snow darkening and suggest that algal pigmentation, which may vary with species identity and physiological state, should be considered alongside biomass when predicting the radiative effects of snow algal blooms.

## IMPORTANCE

Snow algae reduce snow reflectance, increasing solar energy absorption and potentially accelerating snowmelt. Although this effect is commonly linked to algal abundance, snow algae undergo striking changes in pigmentation as cells accumulate the carotenoid astaxanthin. In natural blooms, pigmentation co-varies with biomass, community composition, cell size, and snow physical properties, making its direct optical effect difficult to isolate. Here, we experimentally generated green, orange, and red cyst-like cells from the same snow-derived *Haematococcus* isolate and show that increasing astaxanthin accumulation substantially reduces reflectance, even after accounting for differences in cell abundance and surface area. These results provide direct experimental evidence that pigmentation itself is an intrinsic driver of biological snow darkening. Thus, the optical impact of snow algae depends not only on how much algal biomass is present, but also on its pigment composition, an important consideration for predicting biologically driven snow darkening and melt.

## OBSERVATION

Snow algae blooms are a characteristic feature of melting snow in alpine and polar environments, where they have emerged as important biological drivers of snow darkening (Hotaling et al., 2021). These blooms occur in a striking range of colours, from green to red, reflecting differences in pigment composition associated with taxonomy, life-cycle stage and environmental conditions (Hoham & Remias, 2020). Field studies consistently show that green, orange and red blooms differ in snow reflectance and radiative forcing (Gray et al., 2020; Khan et al., 2021; Almela et al., 2026), while theoretical work predicts that differences in pigment coloration enhances solar energy absorption and melt (Dial et al., 2018). However, the direct contribution of pigmentation to snow reflectance has remained experimentally unresolved because pigmentation co-varies with biological factors (e.g., species composition and biomass) and snow physical properties (e.g., liquid water content and snow microstructure) in natural blooms.

*Sanguina nivaloides* is the predominant species responsible for red snow blooms (Procházková et al., 2019), particularly in high-alpine environments. Despite its ecological importance, no laboratory culture of *S. nivaloides* is available. Although a culture of the related species *S. aurantia* has recently been reported (Raymond et al., 2022), this species produces orange rather than deep red pigmentation, limiting its suitability for investigating the full range of pigmentation associated with predominant *Sanguina* blooms. Consequently, the direct optical effects of algal pigmentation remain experimentally unresolved. To overcome this limitation, we used a *Haematococcus* sp. isolate obtained from a high-alpine snowfield melt pond. The isolate undergoes a reproducible green-to-red pigmentation transition, forming astaxanthin-rich cysts, allowing the optical effects of pigmentation to be isolated experimentally.

Following the experimental approach of Almela et al. (2024), three replicate 24-well plates were incubated for 46 days under combinations of nitrogen and phosphorus availability that generated a gradient of algal pigmentation. Five representative nutrient treatments spanning the orange-to-red pigmentation gradient (samples 1 to 5), together with the corresponding green reference treatment (sample 6; see **Supporting Information**), were selected from each plate and filtered onto 25-mm Whatman GF/F glass microfiber filters. Spectral reflectance (350–800 nm) of the algal biomass was measured from each filter using hyperspectral reflectance spectroscopy prior to pigment extraction.

The experimental treatments yielded cysts spanning a range of pigmentation, from green to orange and red. Astaxanthin concentrations increased progressively from green to orange and red cysts, whereas chlorophyll-*a* concentrations were similar among orange and red samples but significantly lower in green cysts (astaxanthin: F = 90.61, *P < 0*.*0001*; chlorophyll-*a*: F = 46.45, *P < 0*.*0001*; **Table 1**). Consequently, the astaxanthin-to-chlorophyll-a ratio increased progressively from green (0.87 ± 0.06) to orange (1.40 ± 0.48) and red cysts (5.96 ± 0.95) (F = 118.01, *P < 0*.*0001*). The observed increase in astaxanthin is consistent with previous studies showing enhanced carotenoid accumulation under stress conditions, particularly nutrient limitation and high irradiance (Zhekisheva et al., 2002; Leya et al., 2009). In our experiment, the greater biomass achieved under high-phosphorus treatments may have led to more rapid nutrient consumption during the incubation, although nutrient depletion was not directly measured.

**Table 1.** Characteristics of the representative pigmented samples selected for spectral analyses. The table summarizes sample identity (plate and sample), pigmentation category, cell density, total number of cells filtered, astaxanthin and chlorophyll-a concentrations, astaxanthin-to-chlorophyll ratio, and spectral reflectance metrics (area under the reflectance curve [AUC] and mean reflectance). Samples 1–6 correspond to representative pigmentation states generated under different nitrogen-to-phosphorus (NP) nutrient treatments after 46 days of incubation: sample 1, red; samples 2–4, orange; sample 5, light orange; and sample 6, green. For sample 6, spectral reflectance was measured using either the same target cell number as the orange and red samples (6.1) or an increased number of cells to compensate for their smaller size (6.2).

| Plate | Sample | color | Cell density (mL) | Total cells filtered | Astax (ug/mL) | Chla (ug/mL) | Astax:Chla | Reflectance (AUC) | Reflectance (avg) |
| --- | --- | --- | --- | --- | --- | --- | --- | --- | --- |
| 1 | 1 | red | 252,000 | 165,000 | 12.55 | 2.51 | 4.99 | 135.6 | 0.30 |
| 1 | 2 | orange | 274,000 | 165,000 | 1.91 | 1.98 | 0.96 | 161.1 | 0.36 |
| 1 | 3 | orange | 208,000 | 165,000 | 2.79 | 2.31 | 1.21 | 164.4 | 0.37 |
| 1 | 4 | orange | 172,000 | 165,000 | 2.60 | 2.26 | 1.15 | 158.3 | 0.35 |
| 1 | 5 | light orange | 58,000 | 98,600 | 2.93 | 2.48 | 1.18 | 198.3 | 0.44 |
| 2 | 1 | red | 250,000 | 165,000 | 15.89 | 2.65 | 6.01 | 144.3 | 0.32 |
| 2 | 2 | orange | 354,000 | 165,000 | 2.26 | 1.88 | 1.20 | 156.4 | 0.35 |
| 2 | 3 | orange | 274,000 | 165,000 | 2.33 | 1.97 | 1.18 | 155.2 | 0.34 |
| 2 | 4 | orange | 140,000 | 165,000 | 2.30 | 2.12 | 1.08 | 158.0 | 0.35 |
| 2 | 5 | light orange | 36,000 | 61,200 | 2.44 | 2.32 | 1.05 | 197.0 | 0.44 |
| 3 | 1 | red | 212,000 | 165,000 | 18.49 | 2.69 | 6.88 | 121.9 | 0.27 |
| 3 | 2 | orange | 554,000 | 165,000 | 6.45 | 2.70 | 2.39 | 145.7 | 0.32 |
| 3 | 3 | orange | 506,000 | 165,000 | 5.52 | 2.71 | 2.04 | 155.2 | 0.34 |
| 3 | 4 | orange | 182,000 | 165,000 | 3.44 | 2.50 | 1.37 | 136.1 | 0.30 |
| 3 | 5 | light orange | 22,000 | 37,400 | 2.20 | 2.01 | 1.10 | 214.2 | 0.48 |
| 1 | 6.1 | green | 267,000 | 165,000 | 0.57 | 0.62 | 0.91 | 268.2 | 0.60 |
| 2 | 6.1 | green | 358,000 | 165,000 | 0.63 | 0.71 | 0.89 | 281.1 | 0.58 |
| 3 | 6.1 | green | 279,000 | 165,000 | 0.77 | 0.92 | 0.84 | 277.5 | 0.62 |
| 1 | 6.2 | green | 267,000 | 359,700 | 1.22 | 1.44 | 0.85 | 216.6 | 0.48 |
| 2 | 6.2 | green | 358,000 | 359,700 | 0.86 | 0.90 | 0.96 | 223.0 | 0.50 |
| 3 | 6.2 | green | 279,000 | 359,700 | 0.97 | 1.23 | 0.79 | 234.8 | 0.52 |

Spectral reflectance decreased progressively with increasing red coloration of the algal biomass across the visible spectrum, with the largest differences occurring between 400 and 550 nm (**Figure 1**), consistent with the strong absorption of short visible wavelengths by carotenoids. This spectral pattern was consistent across the three experimental replicates. A pronounced absorption feature was also observed near 660 nm, corresponding to chlorophyll-*a* absorption. When reflectance across the full 350– 800 nm range was integrated as the area under the curve (AUC), pigmentation state had a significant effect on reflectance (one-way ANOVA: F = 90.47, *P < 0*.*0001*). AUC declined progressively from green to orange and red cysts. Green reference samples prepared using both equal cell number and surface-adjusted loading exhibited significantly higher AUC values than the red and orange samples, whereas the light-orange sample did not differ significantly from the surface-adjusted green reference.

**Figure 1.**
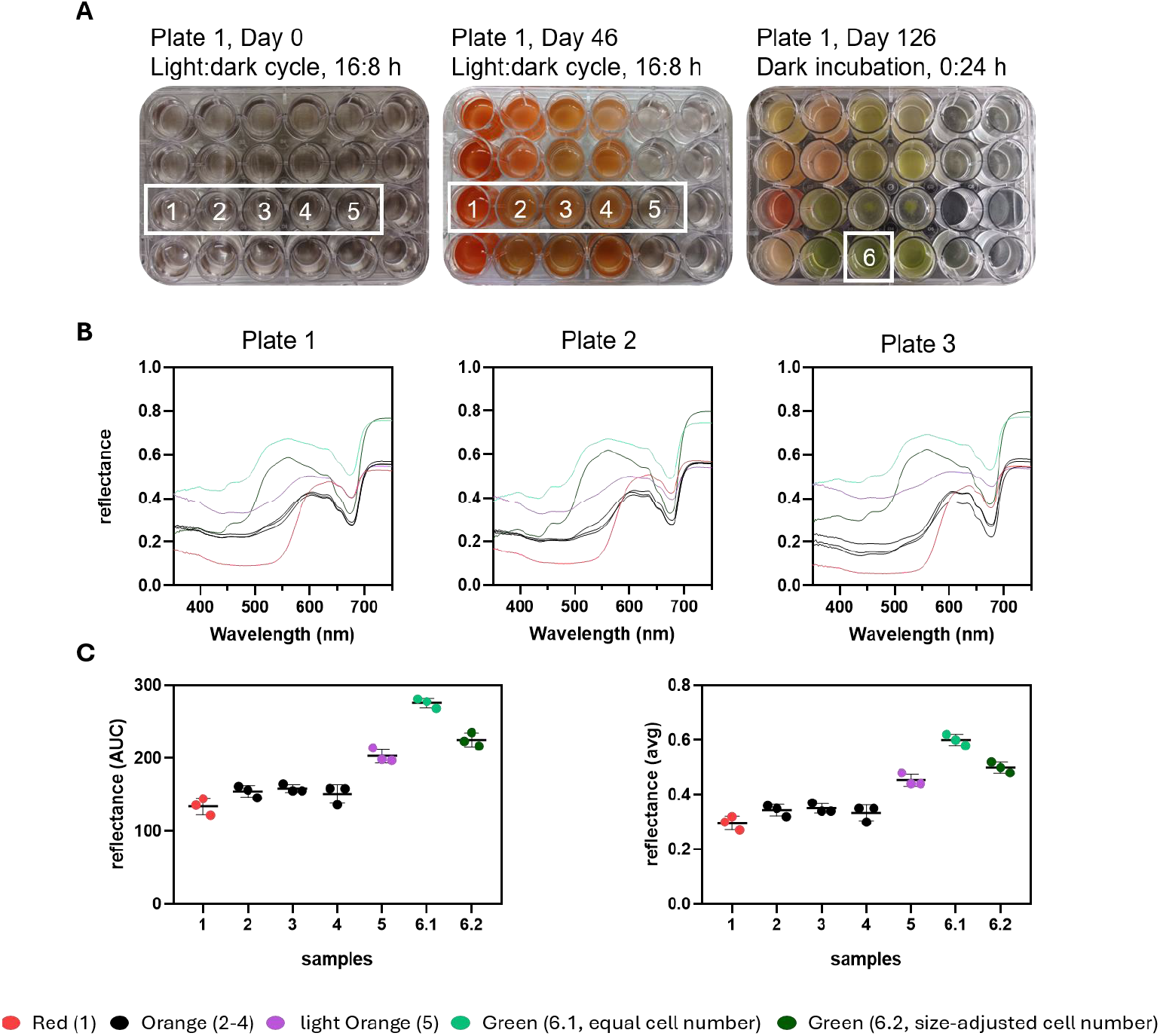
Spectral reflectance of experimentally generated algal pigmentation states. (A) Representative photographs of experimental plate 1 showing the 24 nitrogen–phosphorus (NP) nutrient treatments at the start of the experiment, after 46 days of incubation under a 16 h light : 8 h dark photoperiod, and after a further 100 days of incubation in darkness at 4 °C (day 146), during which some orange samples reverted to green while retaining cyst morphology. (B) Spectral reflectance (350–800 nm) of representative green cyst-like cells (sample 6) containing either the same target cell number as the orange and red samples (light green) or an increased number of cells to account for their smaller size (dark green), together with light orange (sample 5, purple), orange (samples 2–4, black), and red (sample 1, red) cyst-like cells measured from three independent experimental plates. (C) Area under the reflectance spectrum (AUC) and mean reflectance for each sample, showing individual biological replicates and grouped by pigmentation state. Bars represent mean ± SD of the three experimental replicates.

Astaxanthin concentrations were also significantly correlated with AUC (R^2^ = 0.37, *P = 0*.*036*). Consistent with this pattern, the most highly pigmented red cysts (sample 1) exhibited a mean reflectance of 0.30 ± 0.03, compared with 0.60 ± 0.02 for the green reference at equal cell number and 0.50 ± 0.02 for the size-adjusted green reference (sample 6 and 7, respectively). The light-orange sample (sample 5) exhibited an intermediate mean reflectance (0.45 ± 0.02), despite containing fewer cells than the other pigmentation states (**Table 1**). Although biomass could not be standardized for this sample, the result suggests that carotenoid-rich cells (i.e., with an astaxanthin-to-chlorophyll-a ratio > 1) can reduce reflectance more effectively than chlorophyll-dominated green cells, even at lower cell abundances.

Our experimental results help explain why previous field studies have reported contrasting effects of red and green snow on albedo, potentially because differences in snow conditions, particularly liquid water content, confound the optical effects of algal pigmentation (Lutz et al., 2014; Gray et al., 2020; Khan et al., 2021). When present on the snow surface, green snow algae typically occur in wetter snow near snowfield margins, whereas red snow is more often associated with drier, central areas dominated by mature, carotenoid-rich resting cysts. Consistent with our previous field observations of lower reflectance and greater radiative forcing in red than in green blooms (Almela et al., 2026), our controlled experiments indicate that both cell surface area and pigmentation contribute to the optical impact of snow algae. Accounting for differences in effective albedo reduction surface (EARS), which integrates cell abundance and cell surface area (Almela et al., 2024), reduced but did not eliminate the difference between green and red cysts. This supports EARS as a relevant metric of snow algal darkening while identifying astaxanthin accumulation as an additional intrinsic determinant of reflectance.

Greater absorption of solar radiation by astaxanthin-rich cells is expected to increase the energy available for melt, reinforcing positive feedbacks between algal growth and snow ablation (Ganey et al., 2017; Dial et al., 2018). Because both cell size and astaxanthin accumulation can vary with species identity, environmental conditions, and cellular physiological state, the radiative impact of a given cell abundance may vary substantially among blooms and over their development. Biological snow darkening may thus depend not only on how much algal biomass is present, but also on its effective surface area, pigment composition, and distribution within the snowpack (Almela et al., 2025). Incorporating these biological traits into models of snow reflectance may therefore improve predictions of biologically driven snow darkening and melt in alpine snowfields, which constitute an important freshwater resource for human societies.

## ACKNOWLEDGEMENTS

We thank Francesco Caligiore (Thomas D. Niehaus laboratory) for assistance with spectrophotometric measurements. This work was supported by the U.S. National Science Foundation (NSF) through grant DEB-2113784 awarded to T.L.H.

## CONTRIBUTIONS

P.A. conceived the study and designed the experiments. P.A. performed the experiments, analyzed the data, and wrote the first draft of the manuscript. T.L.H. secured funding and supervised the study. Both authors revised and approved the final manuscript.

## COMPETING INTERESTS

The authors declare no competing interests.

## DATA AVAILABILITY

The datasets generated and analyzed during the current study are available from the corresponding author on reasonable request.

## REFERENCES

Almela, P., Elser, J. J., Giersch, J. J., Hotaling, S., Rebbeck, V., & Hamilton, T. L. (2024). Laboratory experiments suggest a limited impact of increased nitrogen deposition on snow algae blooms. Environmental microbiology reports, 16(6), e70052.

Almela, P., Elser, J. J., Giersch, J. J., Hotaling, S., & Hamilton, T. L. (2025). Influence of snow cover on albedo reduction by snow algae. MBio, 16(2), e03630–24.

Almela, P., Elser, J. J., Zmuda, A., Niehaus, T., & Hamilton, T. L. (2026). Community-driven variations in snow algae color modulate snow albedo reduction. New Phytologist, 249(4), 1739–1752.

Dial, R. J., Ganey, G. Q., & Skiles, S. M. (2018). What color should glacier algae be? An ecological role for red carbon in the cryosphere. FEMS Microbiology Ecology, 94(3), fiy007.

Ganey GQ, Loso MG, Burgess AB, Dial RJ. 2017. The role of microbes in snowmelt and radiative forcing on an Alaskan Icefield. Nature Geoscience 10: 754–759.

Gray A, Krolikowski M, Fretwell P, Convey P, Peck LS, Mendelova M, Smith AG, Davey MP. 2020. Remote sensing reveals Antarctic Green snow algae as important terrestrial carbon sink. Nature Communications 11: 2527.

Hoham RW, Remias D. 2020. Snow and glacial algae: a review. Journal of Phycology 56: 264–282.

Hotaling, S., Lutz, S., Dial, R. J., Anesio, A. M., Benning, L. G., Fountain, A. G., … & Hamilton, T. L. (2021). Biological albedo reduction on ice sheets, glaciers, and snowfields. Earth-science reviews, 220, 103728.

Khan AL, Dierssen HM, Scambos TA, Höfer J, Cordero RR. 2021. Spectral characterization, radiative forcing and pigment content of coastal antarctic snow algae: approaches to spectrally discriminate red and green communities and their impact on snowmelt. The Cryosphere 15: 133–148.

Leya, T., Rahn, A., Lütz, C., & Remias, D. (2009). Response of arctic snow and permafrost algae to high light and nitrogen stress by changes in pigment composition and applied aspects for biotechnology. FEMS microbiology ecology, 67(3), 432–443.

Lutz S, Anesio AM, Jorge Villar SE, Benning LG. 2014. Variations of algal communities cause darkening of a Greenland glacier. FEMS Microbiology Ecology 89: 402–414.

Prochazkova, L., Leya, T., Křížková, H., & Nedbalova, L. (2019). Sanguina nivaloides and Sanguina aurantia gen. et spp. nov.(Chlorophyta): the taxonomy, phylogeny, biogeography and ecology of two newly recognised algae causing red and orange snow. FEMS Microbiology Ecology, 95(6), fiz064.

Raymond, B. B., Engstrom, C. B., & Quarmby, L. M. (2022). The underlying green biciliate morphology of the orange snow alga Sanguina aurantia. Current Biology, 32(2), R68–R69.

Zhekisheva, M., Boussiba, S., Khozin-Goldberg, I., Zarka, A., & Cohen, Z. (2002). Accumulation of oleic acid in Haematococcus pluvialis (chlorophyceae) under nitrogen starvation or high light is correlated with that of astaxanthin esters1. Journal of phycology, 38(2), 325–331.

